# Toward a CRISPR-based point-of-care test for tomato brown rugose fruit virus detection

**DOI:** 10.1101/2021.10.29.466394

**Authors:** Joan Miquel Bernabé-Orts, Yolanda Hernando, Miguel A. Aranda

## Abstract

Implementing effective monitoring strategies is fundamental to protect crops from pathogens and to ensure the food supply as the world population continues to grow. This is especially important for emergent plant pathogens such as tomato brown rugose fruit virus (ToBRFV), which overcomes the genetic resistance resources used in tomato breeding against tobamoviruses and has become pandemic in less than a decade. Here we report the development of a CRISPR/Cas12a-based test to detect ToBRFV in the laboratory and potentially in a field setting. Using different tobamoviruses to assess specificity, our test showed a clear positive signal for ToBRFV-infected samples, while no cross-reactivity was observed for closely related viruses. Next, we compared the limit of detection of our CRISPR-based test with a reference real-time quantitative PCR test widely used, revealing similar sensitivities for both tests. Finally, to reduce complexity and achieve field-applicability, we used a fast nucleic acid purification step and compared its results side by side with those of a commonly used column-mediated protocol. The new protocol saved time and resources but at the expense of sensitivity. However, it still may be useful to confirm ToBRFV, detection in samples with incipient symptoms of infection. Although there is room for improvement, to our knowledge this is the first field-compatible CRISPR-based test to detect ToBRFV, which combines isothermal amplification with a simplified nucleic acid extraction protocol.

## INTRODUCTION

Emergent plant viruses are responsible for disease outbreaks that reduce the yield and the quality of many crops (Jones 2021). The occurrence and reach of these outbreaks are accelerating along with globalization and climate change (Trebicki 2020), thus making necessary to implement effective monitoring and eradication strategies.

Tomato brown rugose fruit virus (ToBRFV, genus *Tobamovirus*, family *Virgaviridae*) is an example of an emergent virus that is presently having an important impact on solanaceous crops such as tomato and pepper. ToBRFV has a ssRNA (+) genome encompassing four open reading frames (ORFs) which encode the small subunit of the replicase (RdRp, ORF1), the large subunit of the RdRp (ORF2), the movement protein (MP, ORF3), and the coat protein (CP, ORF4). The genus *Tobamovirus* contains examples of notorious plant viruses including tobacco mosaic virus (TMV) and tomato mosaic virus (ToMV). To date, tomato breeding has taken advantage of the *Tm-1, Tm-2*/*Tm-22* resistance genes to protect new varieties against TMV and ToMV (Pelham 1966, Hall 1980). Nevertheless, ToBRFV overcomes tomato resistance genes (Luria et al. 2017). Furthermore, tobamoviruses are very stable and highly infectious through mechanical transmission (Tomlinson 1987, Smith and Dombrovsky 2020). As a result, ToBRFV has rapidly spread worldwide. Hence, since its first description in the Middle East (Salem et al. 2016, Luria et al. 2017), ToBRFV has expanded to Europe, Asia, and America (Fidan et al. 2019, Ling et al. 2019, Menzel et al. 2019, Skelton et al. 2019, Yan et al. 2019, Alfaro-Fernández et al. 2020, Panno et al. 2020), evidencing the necessity of implementing new measures to prevent its further spread.

Specific testing methods that can be used to detect ToBRFV can critically contribute to the success of intervention and eradication strategies. The most commonly-used method for RNA detection is the reverse transcription real-time quantitative polymerase chain reaction (RT-qPCR). Several protocols have been developed using the RT-qPCR technique to detect ToBRFV, including the International Seed Federation (ISF)-ISHI-Veg protocol (ISF 2020) and others like the set of primers and probe described by Panno et al. 2019. However, RT-qPCR needs to be performed by qualified operators in a laboratory environment, employing equipment that is not always available, and thus limiting its availability especially in low-resource settings. In contrast, isothermal nucleic acid amplification methods such as loop-mediated amplification (LAMP) (Notomi et al. 2000) can be performed over crude plant extracts at a constant temperature by using a water bath or a thermoblock. Some colorimetric LAMP methods have also been developed to easily detect ToBRFV by the naked eye (Sarkes et al. 2020, Rizzo et al. 2021), but these rely on a type of amplification that can be nonspecific. On the other hand, the clustered regularly interspaced short palindromic repeats and its associated proteins (CRISPR/Cas) systems are trespassing the genome editing frontiers and revolutionizing diagnostics. Certain RNA-guided endonucleases such as LbCas12a (former Cpf1) (Zetsche et al. 2015), after recognizing its target DNA, exhibit an unspecific DNase collateral activity *in vitro* that can be exploited to degrade a reporter molecule, thereby adding an extra layer of specificity to the LAMP reaction (Chen et al. 2018) (Figure 1).

**Figure 1:**
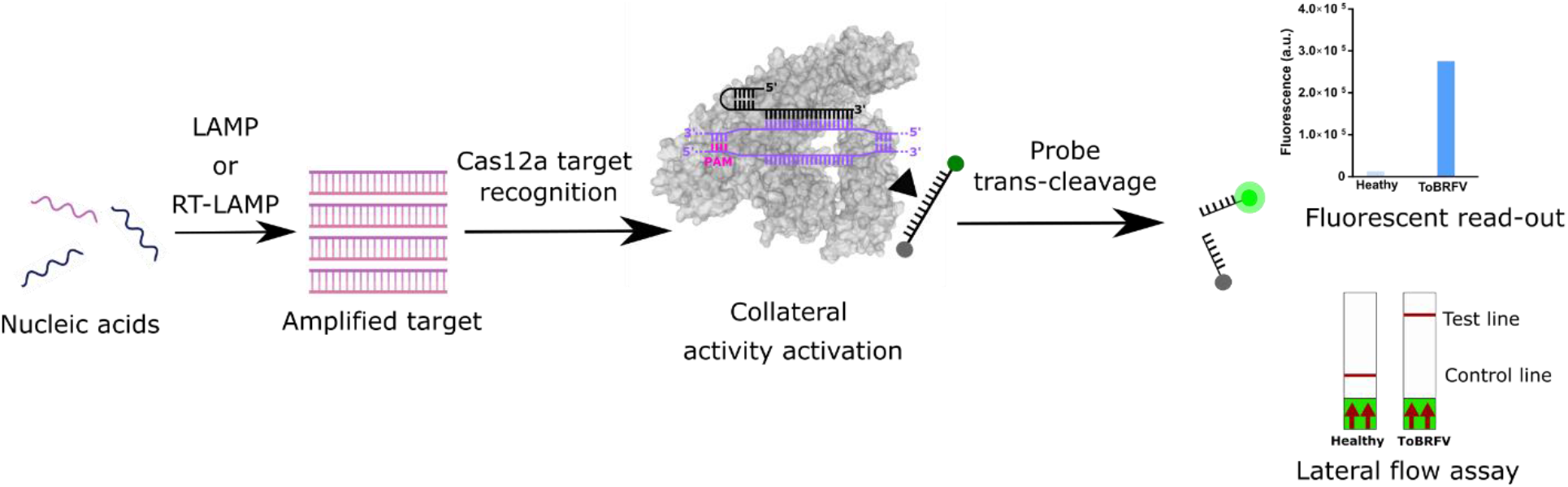
Pipeline for CRISPR/Cas12a-mediated detection processes. A nucleic acid extract is used as a template for the LAMP or RT-LAMP amplification of the target sequences. The amplification product is recognized by the RNA-guided endonuclease LbCas12a through base-pair complementarity of the gRNA and the target sequence, thus triggering the collateral activity of the LbCas12a which, in a non-specific manner, degrades the ssDNA probe revealing the presence of the target sequence. The reporter molecule attached to the probe defines the read-out of the assay, either by fluorescence or using a lateral flow strip.

Typically, a nucleic acid extract is used as a template for the amplification of the targeted sequence by using LAMP (DNA template) or reverse transcription LAMP (RT-LAMP; RNA template). Next, the LAMP product is recognized by the RNA-guided endonuclease LbCas12 through the specific base pairing of the 20-bp target sequence (purple in Figure 1) and the guide RNA (gRNA; black in Figure 1). Target recognition triggers the collateral activity of LbCas12a, producing the unspecific *trans*-cleavage of an ssDNA reporter oligonucleotide. The molecule conjugated to the 3’ end of the ssDNA marks the read-out of the assay. When using a fluorophore, the endonuclease activity dissociates the fluorochrome from the quencher at the 5’ end of the probe, allowing it to be detected. Conversely, the use of a biotin ligand allows the detection of the LbCas12a activity by the naked eye with the aid of a lateral flow assay (LFA) strip (Figure 1). This system has already been employed in the diagnosis of human viruses such severe acute respiratory syndrome coronavirus-2 (SARS-CoV-2) (Broughton et al. 2020, Curti et al. 2021), Influenza A and B (Park et al. 2021), or human papilloma virus (HPV) (Tsou et al. 2019), among others. In plants, isothermal amplification coupled to LbCas12a-mediated detection has also been employed in a handful of studies, mainly for pathogen diagnosis (Aman et al. 2020, Jiao et al. 2021, Mahas et al. 2021) but also for transgenic trait detection (Zhang et al. 2020). Although these works have been carried out to develop field-operative testing methods for pathogen diagnosis (Aman et al. 2020, Jiao et al. 2021, Mahas et al. 2021), only Jiao et al. 2021 actually used nucleic acid isolation protocols that were compatible with field settings. However, this work was carried out using symptomatic samples with a likely elevated titer of the target pathogen, which was perhaps well above the limit of the analytical sensitivity of the detection method. Therefore, the effect of the rapid nucleic acid extraction method on the detection limit of the CRISPR-based tests has not been accounted for and still needs to be evaluated.

Here, we report the design and validation of a CRISPR-based test for the detection of ToBRFV in the laboratory and potentially in a field setting. This test is based on coupling the isothermal amplification of two viral sequences from the ToBRFV movement protein open reading frame, with subsequent detection by using the RNA-guided endonuclease LbCas12a and a specific gRNA. The performance of the CRISPR-based test was compared with a standard RT-qPCR test recommended by the European Plant Protection Organization (EPPO), after which the protocol was adapted for field application.

## MATERIAL AND METHODS

### Virus isolates and plant inoculation

The Spanish ToBRFV isolate from Vicar (Almeria, Spain) (Alfaro-Fernández et al. 2020) was provided by the “Laboratorio de Producción y Sanidad Vegetal” (La Mojonera, Almería, Spain). Isolates of the other tobamoviruses used in this study were acquired from the DSMZ collection: tobacco mosaic virus (TMV, PV-1252), tomato mosaic virus (ToMV, PV-0141), pepper mottle mosaic virus (PMMoV, PV-0093), and tobacco mild green mosaic virus (TMGMV, PV-0124). Approximately 50 mg of dried plant tissue were homogenized from each isolate in 2 mL of 30 mM phosphate buffer pH = 8 using a mortar and pestle. Homogenates were used to mechanically inoculate 25-26 days old leaves (4-5-true leaves) of *N. benthamiana* plants and the first pair of true leaves of 7-10 days-old tomato plants (cultivar M82). For this, the leaves to be inoculated were first dusted with carborundum powder (600 mesh) and then rubbed with the homogenate manually. The plants inoculated were kept separately in a confined greenhouse under controlled conditions set at 16/8 hours photoperiod and 26/22 °C in a day/night cycle. After 10 to 15 days, the leaves that showed obvious symptoms of infection were collected, cut, mixed, and divided into samples of approximately 100 mg. The samples were frozen in liquid nitrogen, ground with a Retsch Mixer Mill MM400 for 1 minute at 30 Hz, and stored at -80 ° C for later analyses.

### Column-mediated RNA extraction

Column RNA extraction was performed using a NucleoSpin RNA plant kit (MACHEREY-NAGEL, Germany) following the manufacturer’s instructions. The RNA was checked by running a 1% agarose gel, its concentration measured with a NanoDrop One (ThermoScientific, USA), and adjusted to a working concentration of 10 ng/µL to be used as a template for the RT-LAMP, RT-PCR, and RT-qPCR.

### RT-LAMP

Loop-mediated isothermal amplification primers (Table S1) were designed using PrimerExplorer v.5 (https://primerexplorer.jp/e/). RT-LAMP reactions were performed using WarmStart LAMP Kit (NEB, USA) at a final volume of 10 μL. Two sets of primers were designed for amplification of the ToBRFV *MP* ORF (MP1 and MP2), and an additional set as a positive detection control (PDC) to amplify the 25S ribosomal RNA from solanaceous species. Primers were added at a final concentration of 0.2 μM for F3 and B3, 1.6 μM for FIP and BIP primers, and 0.8 μM for LF and LB primers. The reactions were performed independently for each set of primers (MP1, MP2, and 25S) using 2 μL of input RNA. The amplification was performed at a constant temperature of 62 °C for 25– 45 minutes (50-90 cycles of 30 seconds each) in an Applied Biosystems StepOnePlus Real-Time PCR System (USA) and tracked with a DNA-intercalant green fluorophore provided with the WarmStart LAMP Kit (NEB, USA). The RT-LAMP amplification was set to 45 minutes for experiments illustrated in Figures 2, 3A and 3B. This incubation was reduced to 25 minutes for experiments in Figures 3C, 3D and 4.

**Figure 2:**
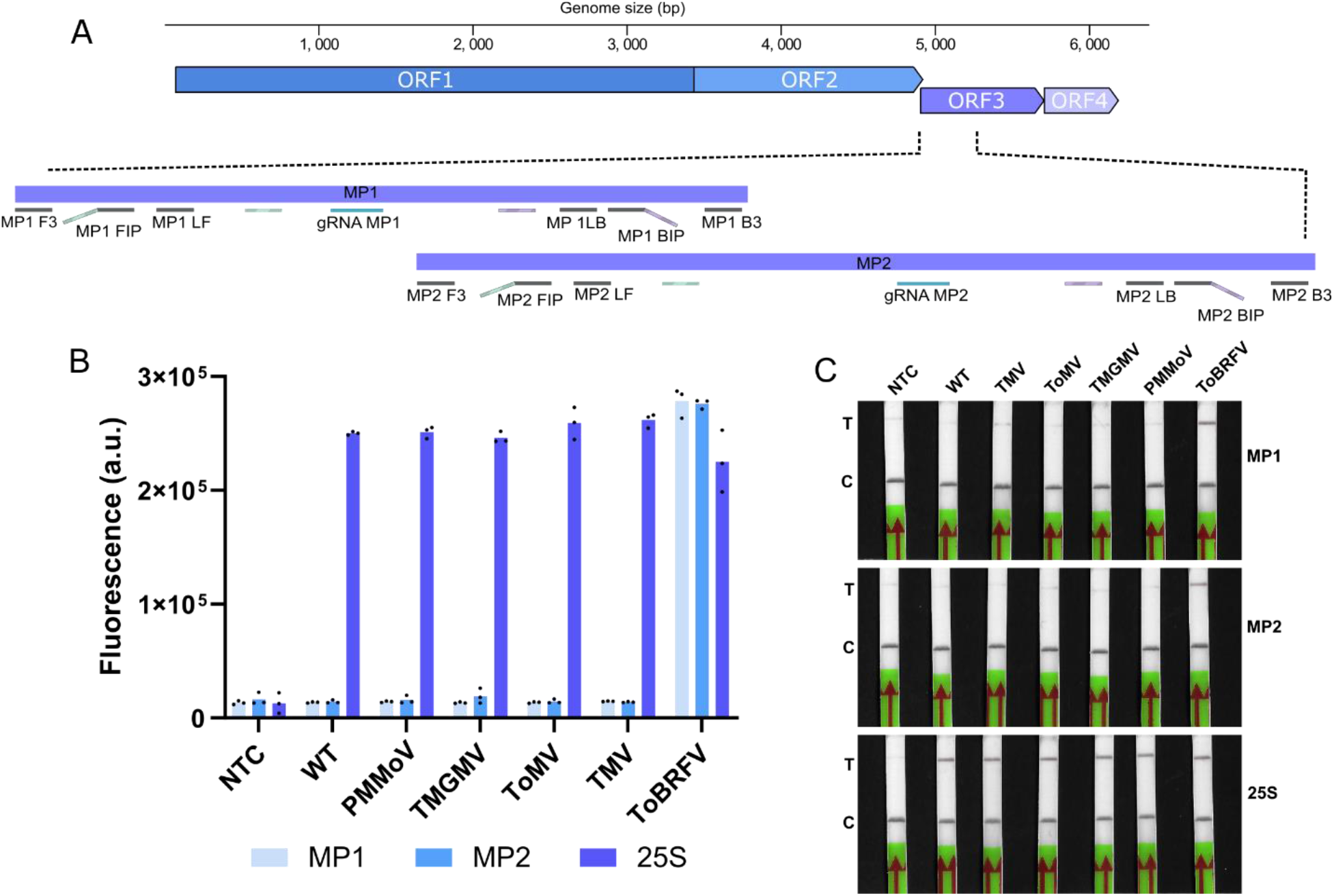
Design and assessment of a CRISPR-based testing method for the detection of ToBRFV *MP* ORF. **A)**. Representation of the ToBRFV genome and the oligonucleotides used for the detection of the *MP* ORF sequence (MP1 and MP2). The position of the RT-LAMP primers is represented by the black rectangles (F3, B3, FIP, BIP, LF and LB). The stripped rectangles represent the binding position of the F1c and the B1c halves of the FIP and BIP primers. The blue rectangles represent the location of the gRNAs (gRNA MP1 and gRNA MP2). An additional set of primers was used to amplify the rRNA 25S as PDC (25S, not shown). **B)** Evaluation of the specificity of the CRISPR-based test MP1, MP2 and 25S targets using a no template control (NTC), a healthy-plant RNA extract (WT) and samples infected with different tobamoviruses (PMMoV, TMGMV, ToMV and TMV) related to ToBRFV, using a fluorescent reporter. Bars represent the average of 3 technical replicates (black dots). **C)** Evaluation of the specificity using a biotinylated reporter with lateral flow strips and TMV-, ToMV-, TMGMV, PMMoV- and ToBRFV-infected tomato plants (T is the test line, and C the control line).

**Figure 3:**
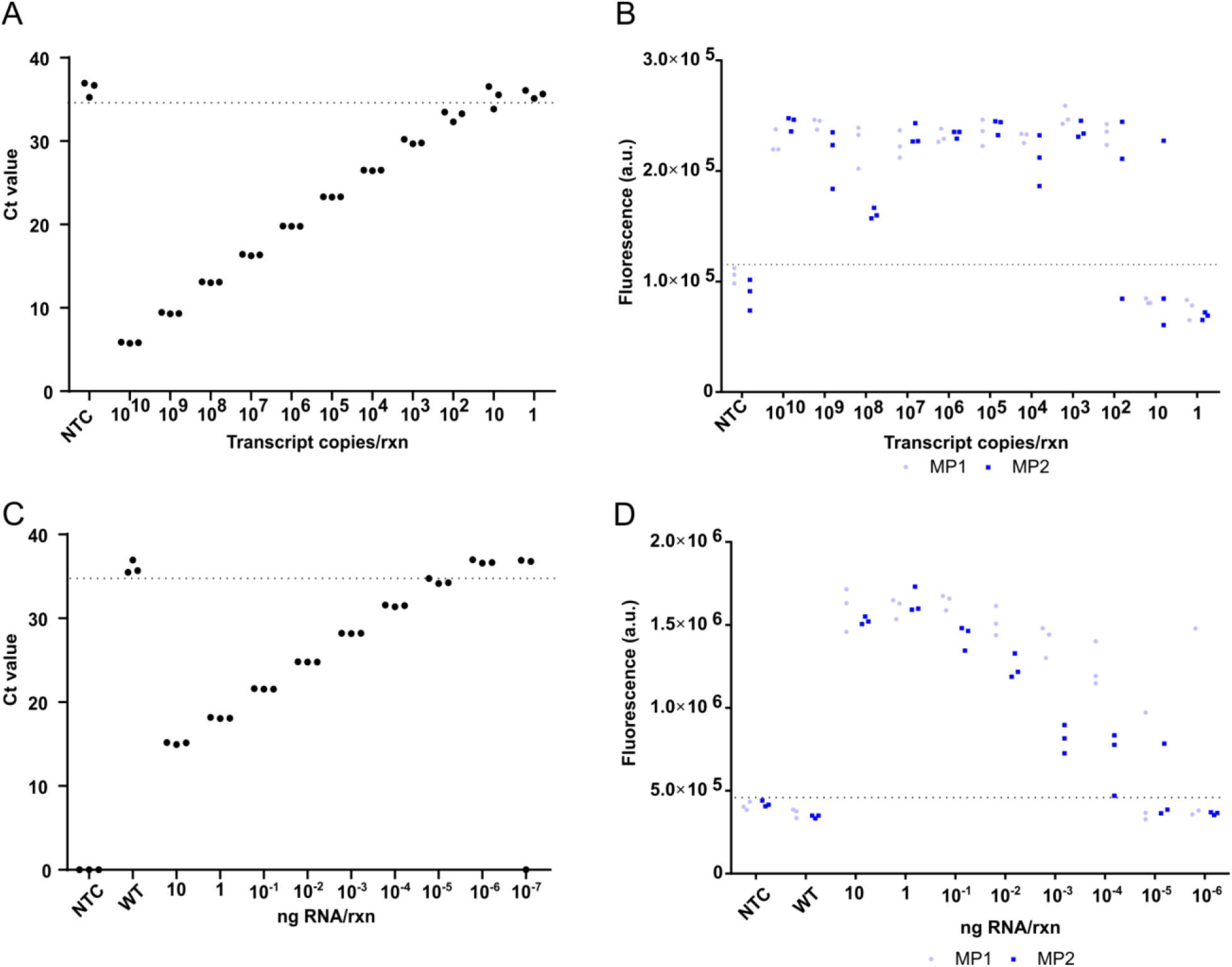
Comparison of the detection limits of the CaTa28 RT-qPCR and the CRISPR-based tests for the detection of ToBRFV. Limit of detection of RT-qPCR and CRISPR estimated with serial dilutions of a synthetic RNA transcript fragment of the *MP* ORF (**A, B**) or an RNA extract from ToBRFV-infected tomato leaves (**C, D**). Each RNA dilution was assessed by triplicate. As negative controls, no template control (NTC) and healthy tomato plant (wild type, WT) RNA extract were used. A healthy tomato plant RNA extract was used as diluent in all cases. The dotted lines show the cut-off value.

### LbCas12a *trans*-cleavage assay

LbCas12a *trans*-cleavage assays were performed using fluorescent and biotinylated reporters, as described by Broughton et al. 2020 and Tsou et al. 2019. When using a fluorescent reporter, 40 nM LbCas12a (EnGen Lba Cas12a, NEB, USA) was pre-incubated with 40 nM of chemically-synthetized gRNA (Synthego, USA, Table S1), and 100 nM of an ssDNA fluorescent (Table S1) reporter for 10 minutes at 37 °C. Then, 2 μL of LAMP product was combined with 18 μL of LbCas12a-gRNA complex and incubated for 10 minutes at 37 °C (20 cycles of 30 seconds each) in an Applied Biosystems StepOnePlus Real-Time PCR System (USA) to detect the resulting fluorescence. When using a biotinylated reporter, the amount of LbCas12a was increased to 100 nM and the gRNA to 125 nm maintaining the pre-incubation time and the 100 nM of the reporter molecule (Table S1). Again, 2 μL of LAMP product was mixed with 18 uL of LbCas12a-gRNA-reporter mixture and incubated for 10 minutes at 37 °C. Finally, 80 uL of 1X HybriDetect assay buffer was added to the reaction mixture, and the lateral flow strip (Milenia HybriDetect - Universal Lateral Flow Assay Kit, Germany) immersed for 2 minutes. Images of the LFA strips were collected using an office scanner and quantified with the ImageJ software.

### RT-PCR

The cDNA was prepared using the Transcriptor First Strand cDNA Synthesis Kit (Roche, Switzerland) following the manufacturer’s instructions, and AB-783 oligonucleotide priming the 3’-UTR of ToBRFV. The resulting cDNA was diluted in 1/10 with water and used as a template for PCR amplification of a fragment of the *MP* ORF, by using Phusion Hot Start II High-Fidelity DNA Polymerase (ThermoFisher, USA) and a final concentration of 0.5 µM AB-782 and AB-783 primers (Table S1). The PCR product was assessed by 1% agarose gel electrophoresis and cleaned using GeneClean Turbo Kit (MP biomedicals, USA) silica gel columns.

### LbCas12a-mediated restriction reaction and efficiency estimation

A restriction reaction was set containing 250 ng of the same PCR product obtained in the previous section “RT-PCR”, 1 µL 1 µM LbCas12a, 1 uL 1 µM gRNA of each target (MP1, MP2 or 25S), 5 µL 10X NEB2.1 and RNase-free water up to 30 µL. The reaction mixture was incubated for 1 hour at 37 °C, after which the restriction product was resolved in a 2% agarose gel for 45 minutes. The efficiency of each target was estimated using ImageJ for the quantification of the digested and nondigested bands. The value of the digested band was divided by the sum of the non-digested plus the digested and the resultant value multiplied by 100 to obtain the %Efficiency shown in Figure S3.

### *In vitro* transcription

RNA fragments of the ToBRFV *MP* ORF were synthesized from a PCR product (see “RT-PCR” section) that included the T7pol promoter. One µL of PCR product was employed to produce the transcripts with the T7 RNA polymerase (Promega, USA), setting a 20 µL-reaction for 2 hours at 37 °C. Then, 3 µL of DNAse I (NEB, USA) were added, and the mixture incubated for 15 minutes at 37 °C. The RNA was precipitated by adding water to 100 µL, 10 µL 3M sodium acetate, 250 µL of ethanol, and incubated for 1 hour at −80°C. The RNA was sedimented by centrifugation at 13,000 rpm for 30 minutes at 4 °C, the supernatant was discarded, and finally, the pellet was air-dried and resuspended with 20 µL of RNase-free water. The resulting RNA was checked as described in the “Column-mediated RNA extraction” section.

### RT-qPCR

Real-time quantitative PCR was performed using KAPA PROBE FAST Universal One-Step qRT-PCR (Roche, Switzerland) and a specific pair of primers and a probe for amplification and detection of ToBRFV (Table S1). Reactions of 10 µL were carried out using a final concentration of 100 mM oligonucleotides and up to 100 ng of RNA template. A StepOnePlus Real-Time PCR System (Applied Biosystems, USA) thermal cycler was employed following this program: reverse transcription at 42 °C for 5 minutes; denaturation at 95 °C for 3 minutes; 40 cycles of amplification with a denaturation step at 95 °C for 3 seconds and annealing and elongation steps at 60 °C for 30 seconds.

### ToBRFV time-course experiment

In total, twelve 5 weeks-old tomato plants cv. M82 (3-4 pairs of leaves approximately) were mechanically inoculated (see above). Three plants per data point (1, 2, 3, 4 days post-inoculation) were sampled, collecting the first pair of new leaves that has been observed to accumulate more virus (van de Vossenberg et al. 2020). Then, the pair of leaves were finely sliced and mixed, after which 100 mg of tissue were introduced into a 2 mL tube with a pair of metal beads for subsequent RNA extraction following the column or the paper strip mediated protocols. These samples were frozen in liquid nitrogen and stored at -80 ° C for later analysis.

### Paper strip-mediated RNA isolation

A rapid RNA extraction protocol was performed following the protocol reported by Zou et al. 2017. Briefly, samples were lysed by shaking them manually in 500 µL of lysis buffer (20 mM Tris pH = 8.0, 25 mM NaCl, 2.5 mM EDTA, 0.05% SDS). Next, a home-made cellulose dipstick was introduced three times in the crude extract to retain the nucleic acids and then washed three times in 1.75 mL of wash buffer (10 mM Tris, pH = 8.0, 0.1% Tween-20). After this, the nucleic acids retained to the dipstick were directly eluted into the LAMP mixture.

## RESULTS

### Design and specificity of a CRISPR-based test for ToBRFV detection

The *MP* ORF (ORF3 in Figure 2A) is the genetic determinant for overcoming the genetic resistance conferred by the *Tm-2*^*2*^ allele to other tobamoviruses such as TMV and ToMV (Hak and Spiegelman 2021); based on this, we reasoned that *MP* may be an appropriate target for our CRISPR-based testing method.

We first screened the ToBRFV *MP* ORF sequence to identify LbCas12a gRNAs candidates (Kim et al. 2017). After filtering candidates considering specificity and structural criteria (Bernabé-Orts et al. 2019), gRNA MP1 and gRNA MP2 were chosen (Figure 2A, Figure S1 and Table S1). Next, we devised two sets of RT-LAMP primers flanking each gRNA target (Figure 2A, Figure S1 and Table S1). From this point on, we will refer to each set of gRNA and primers as MP1 and MP2, respectively. Importantly, MP1 and MP2 overlap the ISF-ISHI-veg RT-qPCR CaTa28 test (ISF 2020), facilitating comparisons. We also included a complementary PDC named 25S, which targets the 25S subunit of the ribosomal RNA from solanaceous species. Using RNA extracts obtained from plants infected with different tobamoviruses (PMMoV, TMGMV, ToMV, TMV and ToBRFV) we demonstrated that our CRISPR-based test can detect ToBRFV without cross-reactivity with closely related viruses (Figure 2B). While all of the samples were positive for the 25S PDC, only the ToBRFV-infected sample gave rise to fluorescent signals above the background levels for all the three targets, indicating that ToBRFV-negative results were due to the absence of this virus rather than a testing failure of MP1 or MP2. This result was corroborated when using LFA strips with the TMV-, ToMV-, TMGMV-, PMMoV- and the ToBRFV-infected samples (Figure 2C). The ToBRFV extract showed intense test lines for MP1, MP2 and 25S, while the rest of samples were positive only for the 25S target, aligning with the previous results obtained with the fluorescent reporter. Interestingly, the test lines of the MP2 and the 25S targets were more intense than that for the MP1. An evaluation of the efficiency of the three gRNAs showed that MP2 gRNA was the most efficient (62%), followed by the 25S (48%) and the MP1 (39%) (Figure S2), thus revealing a correlation between the gRNA efficiency and the LFA output as Zhang et al. (2020 previously showed. Altogether, these results confirmed that our CRISPR-based testing method specifically detected ToBRFV while discarding false negatives at the same time, thanks to the testing process with the 25S PDC. Importantly, the test also allowed the adaptation of the visualization of the results by fluorescence and/or by the naked eye.

### Performance of the CRISPR-based test for ToBRFV detection

Next, we sought to assess the performance of our CRISPR-based detection platform and compare it with the CaTa28 RT-qPCR test described in the ISF-ISHI-Veg protocol (ISF 2020), supported by the EPPO (Figure 3). This RT-qPCR test is directed to the *MP* ORF of ToBRFV, near our MP1 and MP2 target sites, thus ruling out a possible bias in the efficiency due to positional effects. First, the limit of detection of both methods was assessed using serial dilutions of a synthetic transcript encoding a fragment of the *MP*, to accurately evaluate the number of viral copies per reaction (rxn) that each method was able to detect (Figure 3A and 3B). These experiments revealed that under our particular conditions of equipment, material and reagents, the CaTa28 RT-qPCR test reliably detected up to 100 copies of transcript/rxn, although one technical replicate corresponding to the 10 copies/rxn dilution was also positive (Figure 3A). Similarly, our CRISPR-based method detected up to 100 copies/rxn dilution, although with some difficulties for the MP2 target, which failed to detect the synthetic transcript in one technical replicate (Figure 3B). The sensitivity of both techniques was also compared using serial dilutions of an RNA extract obtained from a tomato plant infected with ToBRFV (Figure 3C and 3D). In this case, the CaTa28 RT-qPCR test (Figure 3C) outperformed the CRISPR-based tests (Figure 3D) by detecting the viral RNA in a dilution one order of magnitude higher (10^−5^ ng/rxn) than the CRISPR method (10^−4^ ng/rxn). Although some technical replicates were positive beyond this 10^−4^ ng/rxn dilution, this was the last point consistently detected, suggesting that this is the limit of detection of our CRISPR-based method. In summary, these results showed that our CRISPR detection method possesses a similar sensitivity to the CaTa28 RT-qPCR, at least when a synthetic transcript is used as a template. When using dilutions of an RNA extract, the CRISPR test showed one order of magnitude less sensitivity than the RT-qPCR test.

### Development of a rapid protocol for CRISPR testing

RNA isolation is a critical step, which consumes most of the time and resources of the testing process, influencing at the same time the output of the subsequent detection method. For this reason, we sought to couple a rapid RNA extraction protocol, which requires a paper strip to retain the nucleic acids (Zouet al. 2017), with our CRISPR-mediated testing protocol and compare it with the standard column-mediated RNA extraction protocol that we had been using in our previous experiments (Figure 2 and 3). To this end, we generated a set of samples by mechanically inoculating twelve tomato plants (P1-P12) and sampling them at different days post-inoculation (dpi) in subsets of three biological replicates. Our purpose was to obtain a collection of samples with a variable range of viral load, use them to extract the genetic material of the virus with both protocols and compare the performance of our test after that.

Figure 4A shows the data obtained when using the column-mediated RNA extraction protocol. These results revealed that ToBRFV could be detected in systemically-infected tissue as soon as 1 dpi, at least for plant 1 (P1) which was positive to both MP1 and MP2 targets. P2 and P3 were positive only for some replicates and only for one target. On the following time points, only P6 was negative, detecting one replicate of the MP2 target. ToBRFV negative plants were positive for the 25S PDC, indicating that the absence of signal for MP1 and/or MP2 targets reflected a low titer or absence of the virus in these samples, rather than test failure. Overall, the signal intensity of the MP1 and MP2 targets increased as the infection progressed, indicating an increase in the viral load. For example, at 1 dpi, the signal of P1 for the MP1 and MP2 targets was 3-4 times higher, respectively, as compared to the WT sample, whereas this difference was 10-14 times higher at 4 dpi. In contrast, the signal intensity of the 25S target remained steady through all the data points, indicating the stability of the RNA extraction for all the time points. When using the paper strip-mediated RNA extraction protocol (Figure 4B), clear positives were only detected at 4 dpi (P10-P12). Before this time point, some biological replicates were inconsistently positive (e.g. P1 or P8), not allowing for a clear diagnosis of these samples with this protocol. Furthermore, in general, the signal intensities were lower than for column-mediated RNA extractions (compare Figures 4B and 4A). Finally, to check the visual output using the LFA strips, we selected the replicates that were positive in the previous experiments (Figure 4A and 4B) and used them to perform the LbCas12a-mediated detection, this time with a biotinylated reporter. The signal intensities of the LFA strips test lines shown in Figure S3 were quantified, and the resulting values plotted in the heat-maps shown in Figure 4C (column-mediated extraction) and 4D (paper strip extraction). As expected, the results aligned perfectly well with those obtained with the fluorescent reporter. Even the replicates that displayed low fluorescent signals close to the detection limit (e.g. MP2 target on P1 Figure 4A; or MP1 target on P1 in Figure 4B) were easily detected with the LFA approach, confirming that fluorescent results can be extrapolated to naked eye visualization. In sum, despite the considerable reduction in time, labor, reagents, and equipment associated with the paper strip-extraction procedure, the column-mediated protocol was the most efficient, allowing detection of ToBRFV systemic infection 3 days earlier than the paper strip protocol.

**Figure 4:**
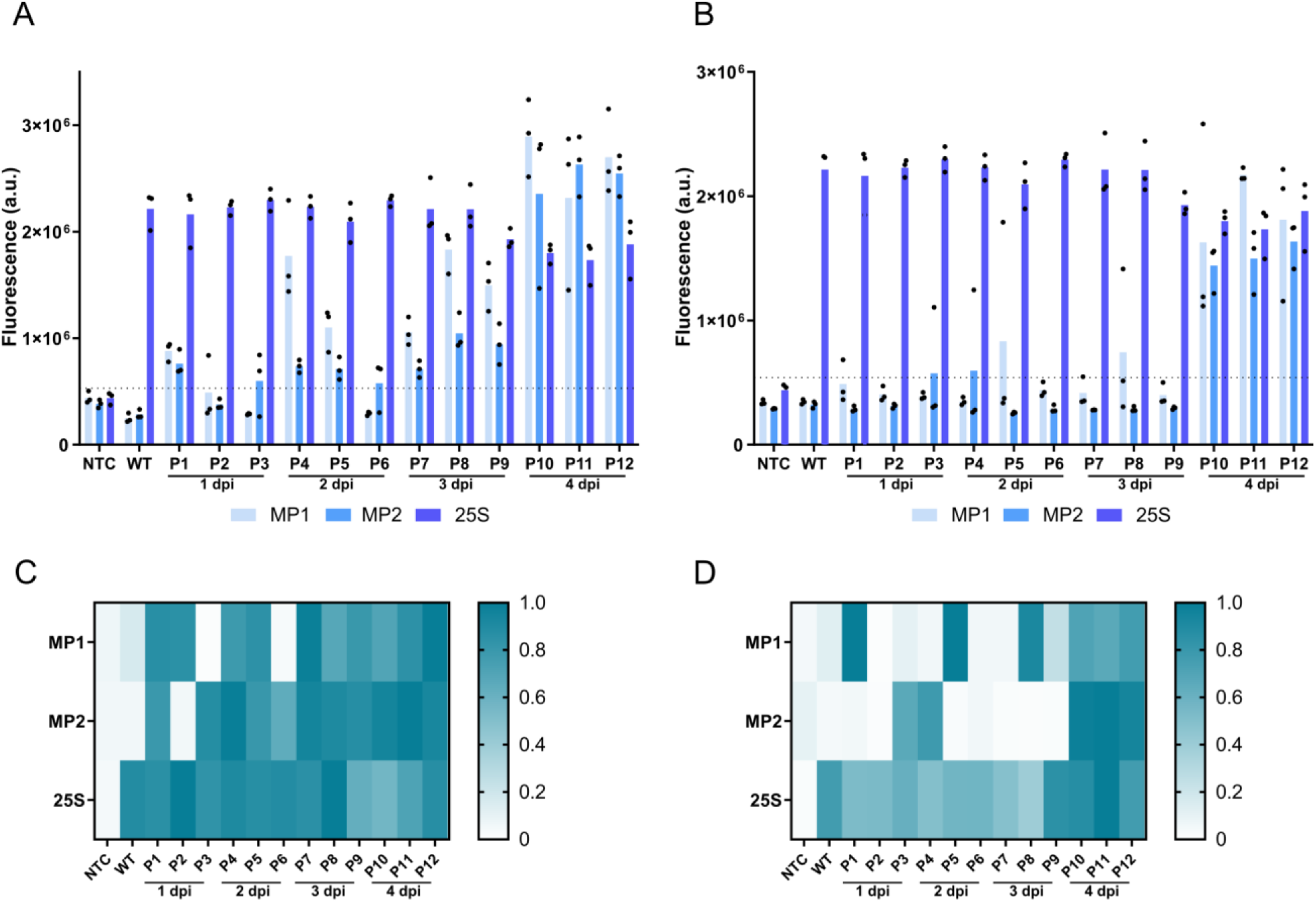
Influence of the RNA extraction protocol on the CRISPR-based detection. Twelve ToBRFV-inoculated tomato plants (P1-P12) were sampled at 1, 2, 3 and 4 days post-inoculation (dpi), three plants each time. The pair of nascent leaves were collected and the sampled tissue was finely sliced, mixed and split into two subsamples used for the conventional column-mediated RNA extraction protocol, or a rapid paper strip-mediated nucleic acid extraction protocol to capture the nucleic acids (Zou et al. 2017). **A, B)** Fluorescent read-out from the column-extracted samples (**A**) and the rapid-extracted samples (**B**). Bars represent the average of three replicates (black dots). **C, D)** Signal quantification from the LFA strips of the column-extracted samples (**C**) and the rapid-extracted samples (**D**). Each value was normalized to the highest value. As negative controls, no template control (NTC) and healthy tomato plant (wild type, WT) RNA extract were used. Quantification of the test line was performed using ImageJ. The dotted lines in A and B show the cut-off value.

## DISCUSSION

One important step in disease control is the unambiguous identification of the pathogen to limit or prevent its spreading. RT-qPCR tests have become the gold standard due to their versatility and ability to detect a few molecules of nucleic acid from the disease agent. However, the technical complexity of RT-qPCR restricts its application to laboratory settings.

CRISPR-based diagnostics promises to transfer the versatility, sensibility, and specificity of genetic testing methods from the laboratory to field settings, accelerating the decision-making process while reducing costs. In the past few years, the natural diversity of CRISPR/Cas systems has been harnessed to develop many testing methods with different chemistry and analyte specificity. For a sensitive detection, all of these methods require a pre-amplification step of the targeted sequence from the pathogen genome. The difference among these methods stems on the molecule that the Cas enzyme can recognize. Thereby, the endoribonuclease Cas13a recognizes a ssRNA analyte (Gootenberg et al. 2017), the endonuclease Cas12a a dsDNA (Chen et al. 2018), and the Cas14 a ssDNA (Harrington et al. 2018). This directly influences the chemistry of the detecting process, imposing some limitations. For example, when using the Cas13a system, an RNA polymerase is needed to transcribe the amplified dsDNA target sequence to a ssRNA moiety which can be recognized by the endoribonuclease. Similarly, Cas14a requires a T7 exonuclease to obtain a ssDNA sequence from the amplified target. Conversely, the LbCas12a can detect the dsDNA amplification product directly, saving steps that require additional enzymatic activities. Therefore, to our understanding, we selected the most straightforward approach, using the LbCas12a endonuclease to develop our CRISPR-based testing method for ToBRFV.

A pre-amplification step needs to be carried out to improve the sensitivity of CRISPR-mediated detection method. Usually, to circumvent the utilization of sophisticated thermal-cyclers, isothermal amplification methods such as recombinase polymerase amplification (RPA) (Piepenburg et al. 2006) or LAMP (Notomi et al. 2000), are the best choice for point-of-care applications. Both isothermal amplification systems are comparable in terms of sensitivity, rate of amplification, and tolerance to inhibitors (Zou et al. 2020). The main differences are found in the enzymatic activities involved in the process, which determines the molecular mechanism underlying the amplification and the temperature of incubation. Thus, similar to PCR, RPA only requires two oligonucleotides and can work at 37-42 °C. In contrast, the LAMP reaction involves six oligonucleotides and higher incubation temperatures of 60-65 °C. Apparently, as RPA and LbCas12a share the same incubation temperature of 37 °C, this may be the best match for a single-tube amplification plus detection reaction. However, the different chemistry of both processes complicates their combination into a single-tube reaction, or at least reduces the sensitivity of the detection as described by Kellner et al. 2020 when using Cas13a. Mahfouz’s group used this approximation for plant virus detection, although no direct comparison with other testing methods was carried out (Aman et al. 2020). In addition, the simplicity of the RPA is a double-edged sword, as this technique is more tolerant to mismatches and thus to spurious amplifications (Li et al. 2020). Altogether, we decided to use LAMP and LbCas12a in a two-tube reaction and eventually upgrade our protocol to a single-tube reaction. To this end, the amplification and detection reactions can be physically separated with an inorganic phase, as previously done by Wang et al. 2021, or combined by using a thermostable Cas enzyme such as AapCas12b endonuclease (Joung et al. 2020). In this latter work, an extensive screening of chemical adjuvants was conducted to enhance the simultaneous amplification and detection reactions, evidencing the incompatibility of both processes, and reinforcing our strategy.

Hence, we targeted two different positions of the *MP* ORF (MP1 and MP2) and added a PDC (25S), thus facilitating the interpretation of the results. Our ToBRFV CRISPR-based assay was considered positive if there was detection in MP1 and MP2, or presumptive positive if there was detection in either MP1 or MP2. A negative test would be when only 25S was detected. Examples of these outputs can be found in Figure 4. First, we confirmed the specificity of our test using plants infected with different tobamoviruses, showing that only ToBRFV-infected samples gave rise to a strong signal for both MP1 and MP2, whereas the 25S PDC was detected in all the samples except NTC. Next, we assessed the detection limit of our CRISPR-based test by comparing it with the CaTa28 RT-qPCR test standard. Note that to capture all the amplification events, including samples with a low number of copies, the amplification time was set to 45 minutes in a first instance (Figure 3A and 3B). However, being aware that a long amplification time might reduce the field-applicability of our CRISPR-based method, this incubation was reduced to 25 minutes in the following experiments, illustrated in Figures 3C, 3D and Figure 4. Using a synthetic RNA extract as a template, the CaTa28 test and our CRISPR-based test showed equivalent detection limits. Conversely, when using dilutions of an RNA extract obtained from a ToBRFV-infected tomato plant, the CaTa28 RT-qPCR test showed a detection limit one order of magnitude higher than our CRISPR-based test. Furthermore, the signal from the CRISPR test decreased with the magnitude of the RNA dilutions, which may reflect a decrease in the LAMP product due to the reduction in the amplification time. Altogether, these results showed that our CRISPR detecting method possesses a similar sensitivity than the CaTa28 RT-qPCR test, at least when the LAMP amplification time was extended, and a synthetic transcript used as a template. Decreasing to half the amplification time reduced the limit of detection one order of magnitude, but shortened the detection process considerably. Finally, to adapt our test to a field setting, we shifted from a conventional column-mediated RNA extraction protocol to a rapid RNA extraction protocol which only involves a strip of paper to adsorb the nucleic acids present in a crude plant extract (Zou et al. 2017), as previously used by Zhang et al. 2020 to detect the *Bt* transgene in rice. The results revealed that the RNA extraction protocol critically influenced the output of the test. The column-mediated protocol outperformed the rapid one, which was only able to detect ToBRFV at 4 dpi, when the plants presented slight symptoms of infection such as subtle blistering and/or mottling. Therefore, in practice, this rapid protocol could be reserved for plants which present clear symptoms of virus infection. In the future, more rapid extraction procedures such as that described by Silva et al. 2018 need to be tested in order to assess its compatibility and influence on the performance of our assay.

While writing this manuscript, Ziv Spiegelman’s group reported on the use of CRISPR/Cas12a to detect ToBRFV (Alon et al. 2021). Notably, the authors designed and screened efficient gRNAs for the differential detection of ToMV and ToBRFV, and evaluated their ability to detect serial dilutions of an end point PCR product using a fluorescent reporter. In spite of the significance of the work, important aspects for virus detection such as an extensive validation of the assay specificity by using several tobamoviruses, or the estimation of the detection limit, were not addressed in this study. Our work was aimed at developing a CRISPR-based test to detect ToBRFV, which may potentially be deployed to the field. To this end, we contemplated all the steps involved in the testing process, including the amplification of the RNA template, and not only the CRISPR/Cas12a-mediated detection. We chose an isothermal amplification method such as RT-LAMP which is compatible with a field setting, in comparison with the RT-PCR used by Alon et al. 2021. Importantly, the isothermal amplification imposes some design constraints (e.g., primers, amplicon size) that drastically reduces the target selection, so a good gRNA may not necessarily match with a good primer set for amplification. Therefore, the selection of a proper primer set in conjunction with the gRNA design is crucial for achieving good sensitivity levels, as Joung et al. 2020 demonstrated. In addition to a field-compatible amplification method, we also checked the sensitivity of our CRISPR-based test coupled to a paper strip-mediated RNA isolation protocol, which is also applicable outside of the laboratory. Finally, aside from the fluorescent reporter, we also used LFA strips which are usable in the field. Altogether, we believe that our test is closer to field deployment for the following reasons: (i) it can be coupled to rapid nucleic acid extraction, (ii) it is based on isothermal amplification, (iii) its results can be read by the naked eye using LFA.To our knowledge, this is the first report of a CRISPR-based test to detect ToBRFV developed to be as close as possible to field deployment. Although its applicability under field conditions is still debatable, this work paves the way for further improvements which may soon lead to simple and effective tests for the detection of ToBRFV in the field.

## ACKNOWLEDGMENTS

We thank Mario Fon for editing the manuscript and Mari Carmen Montesinos and Julia Muñoz for technical support.

## AUTHOR CONTRIBUTIONS STATEMENT

J.M.B.O., Y.H. and M.A.A. conceived the research. J.M.B.O. and M.A.A. designed experiments. All the experiments were done by J.M.B.O., who also wrote the manuscript under the supervision of Y.H. and M.A.A. All authors read and approved the final manuscript.

## ADDITIONAL STATEMENTS

1. No plant or seed specimens have been collected for this work.
2. No voucher specimens have been used. Virus isolates are deposited and are publicly available from the Leibniz Institute-German Collection of Microorganisms and Cell Cultures GmbH (https://www.dsmz.de/dsmz).

## SUPPLEMENTARY FIGURES

**Figure S1:**
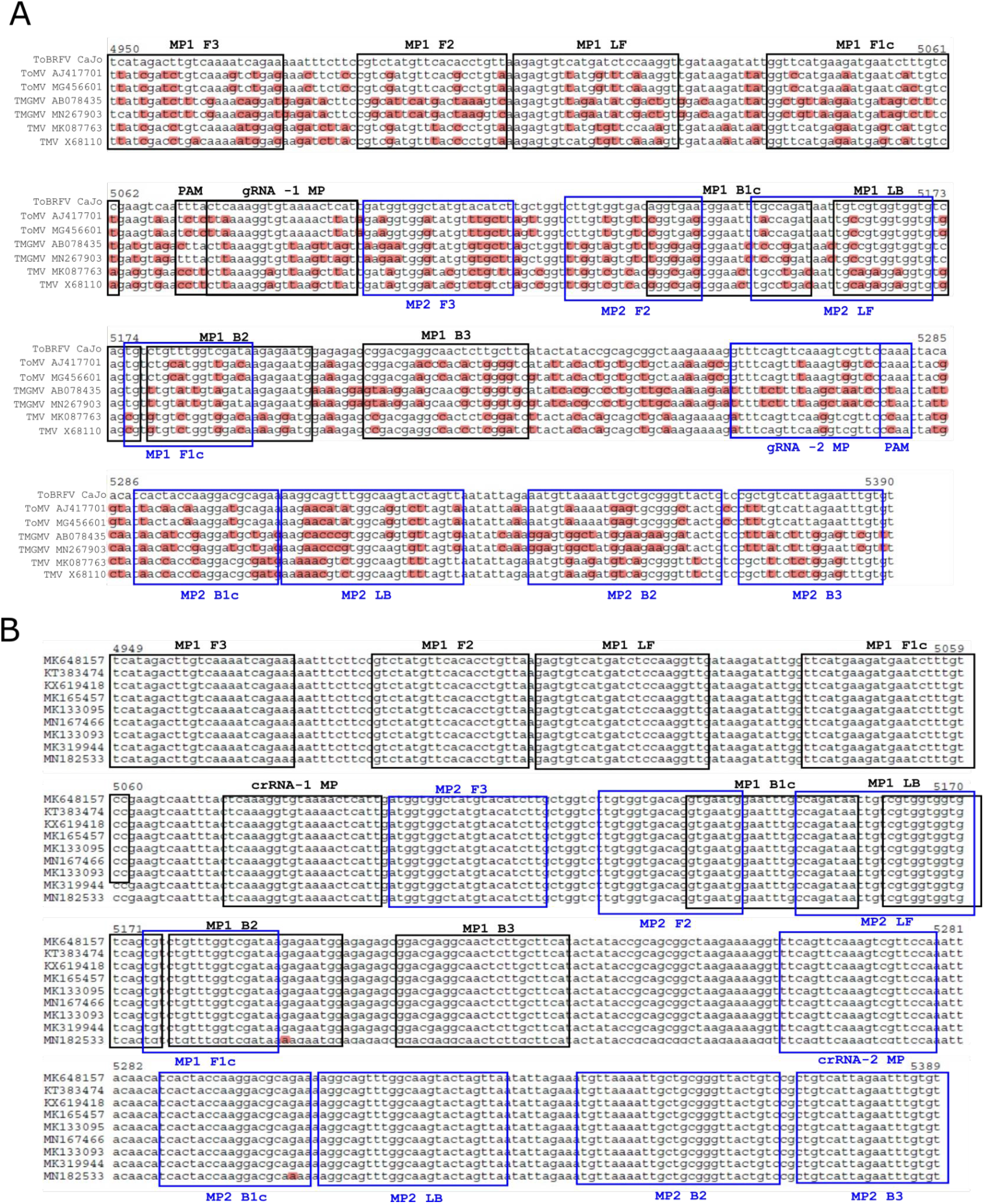
*In silico* evaluation of the exclusivity (A) and inclusivity (B) of the oligonucleotides designed for the CRISPR-mediated ToBRFV detection. Only ToBRFV accessions are shown in (B). The oligonucleotides for the amplification and the detection of the *MP* ORF are highlighted in black for MP1, or in blue for MP2. The sequences of the viruses were retrieved from the NCBI nucleotide database. The accession number is shown together with the acronym of the virus. Sequence alignments were performed using ClustalW and Benchling webtool.

**Figure S2:**
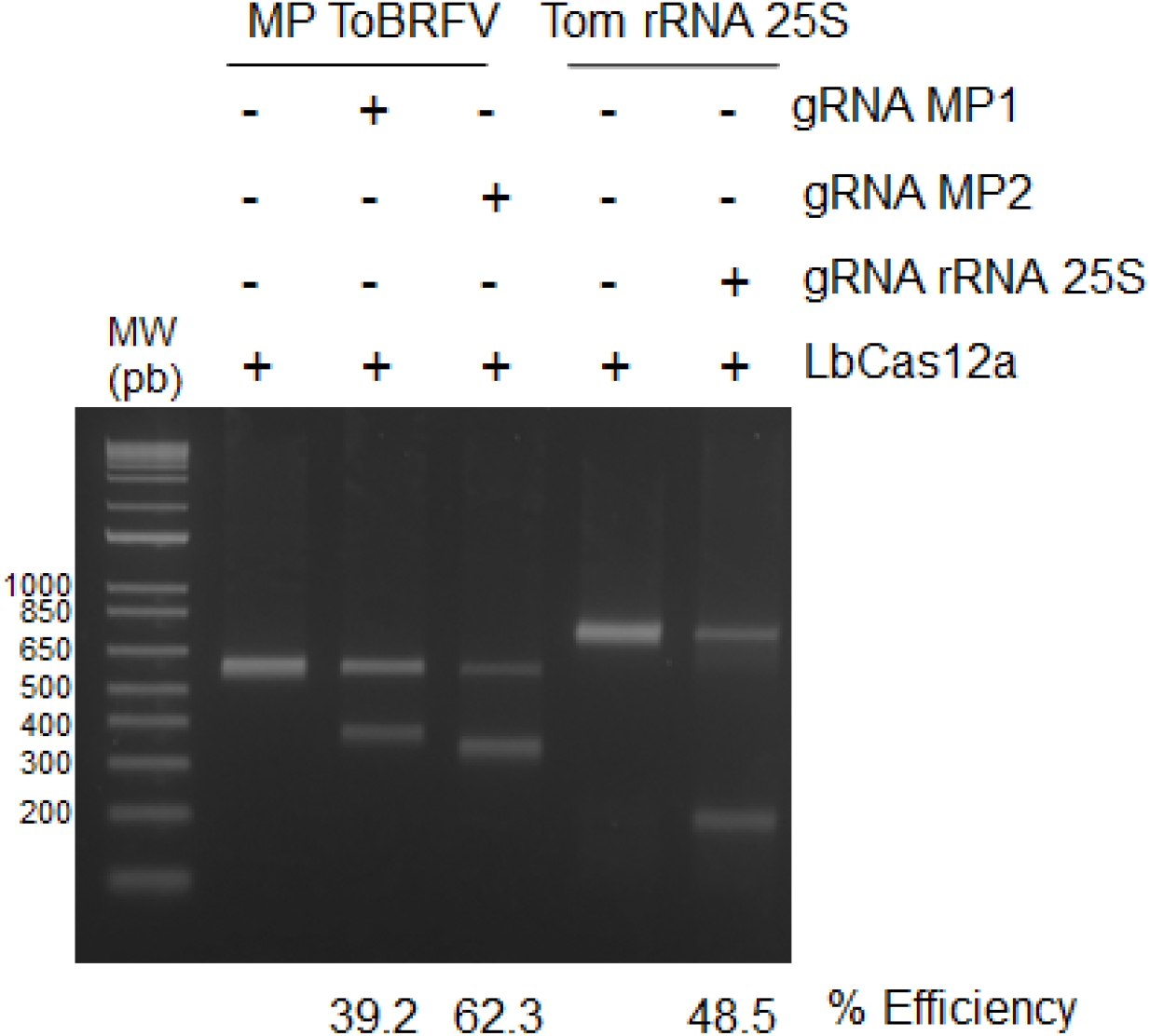
Assessment of MP1 and MP2 gRNAs efficiency through LbCas12a-mediated restriction of PCR products containing the targeted sequence and subsequent electrophoresis in a 2% agarose gel.

**Figure S3:**
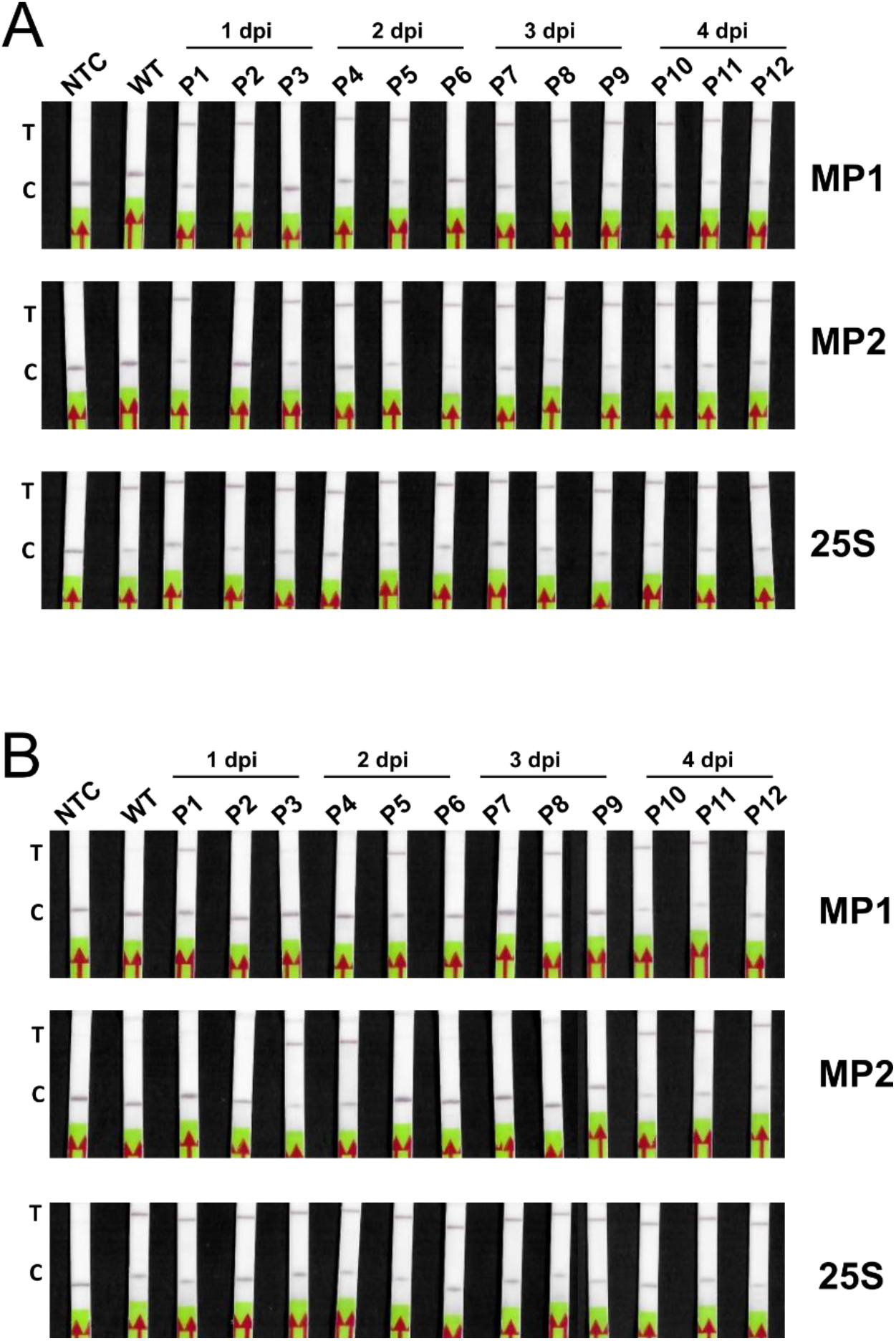
Lateral flow strips derived from the column-mediated RNA extraction protocol (A) and the rapid paper strip-mediated protocol (B). The signal in these strips was quantified using ImageJ, and the results used to represent the heat-maps shown in Figures 3C and 3D.

## SUPPLEMENTARY TABLE

**Table S1:**
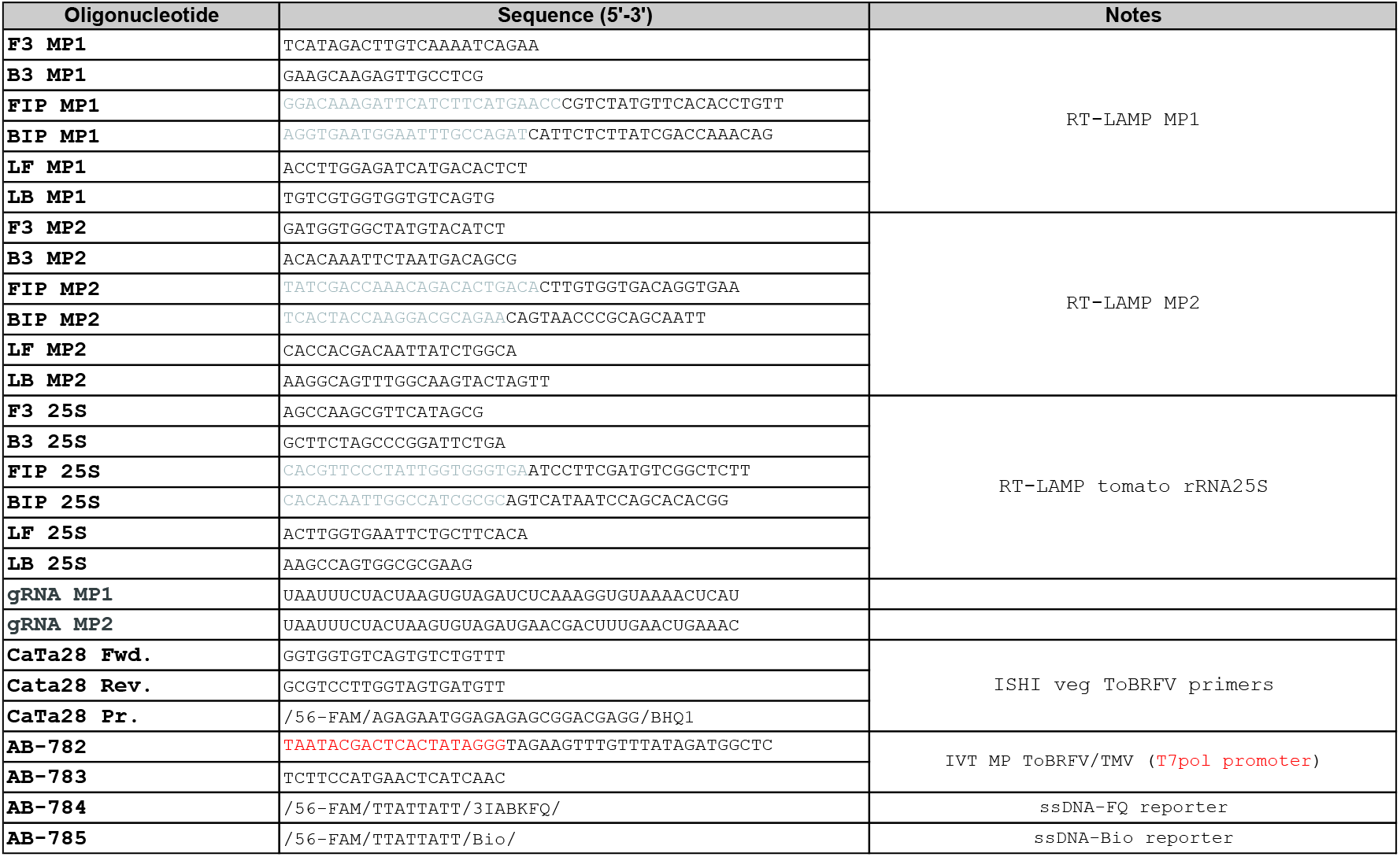
oligonucleotides used in this study.

